# Cell-specific transcription dysregulation in human Huntington’s disease-positive developing striatum

**DOI:** 10.64898/2026.06.19.733377

**Authors:** Sophie V. Precious, Oliver J.M. Bartley, Phoebe Linehan, Alys N. Aston, Rachel Hills, Anne-Marie McGorrian, Vincent Dion, Anne E. Rosser

## Abstract

Huntington’s disease (HD) is an autosomal dominant neurodegenerative disorder caused by a CAG-repeat expansion in the *HTT* gene. Progressive loss of striatal projection neurons leads to cognitive, psychiatric, and motor impairments that typically manifest in midlife, despite the presence of the expansion from conception. Increasing evidence supports a neurodevelopmental component to HD; however, authentic human developing HD striatal tissue has not previously been characterised.

We analysed an HD positive human fetal striatal sample alongside an age- and sex-matched control. CAG-repeat length was determined, and single-cell RNA sequencing was used to investigate gene expression. We compared the fetal HD transcriptional signature with publicly available datasets from postmortem adult HD brain tissue.

We identified 2,032 differentially expressed genes and defined nine cellular clusters, each exhibiting distinct transcriptional profiles. Gene enrichment analysis revealed disruption of key biological processes across the developing HD striatum, with pathway-level dysregulation varying between clusters. There was overlap in gene expression changes between fetal and adult HD striatal tissues.

Together, these findings demonstrate that molecular features of HD pathology are present during early human striatal development, supporting the concept that disease mechanisms are established decades prior to clinical onset.

## Introduction

Huntington’s disease (HD) is an inherited, autosomal dominant, neurodegenerative condition, caused by an expanded CAG-repeat (>36) in the *HTT* gene; characterised by progressive psychiatric, cognitive and motor dysfunction. The disease-causing mutation (*mHTT*) is present from conception, but clinical deficits typically only appear in adulthood.^1^

Although *HTT* is ubiquitously expressed, the main organ affected in HD is the brain, and the predominant cells affected are the medium spiny projection neurons (MSNs) of the striatum. The earliest structural change is a small reduction in striatal volume (specifically, the putamen) ∼24 years before clinical onset; with parallel elevation of biomarkers (neurofilament light and *YKL*-40) in CSF, particularly in individuals closer to predicted onset.^2^ Somatic CAG-repeat expansion also predates disease onset, and somatic expansion scores in blood DNA predict age-of-onset in HD.^3^

Together, these findings demonstrate that the biological impact of *mHTT* is felt decades prior to clinical onset. However, as the gene is present from conception, the question arises as to how early this occurs. There is mounting evidence to suggest a developmental impact of *mHTT*; studies in rodents and human fetal cortical cells demonstrate that *mHTT* affects brain development.^4–7^ There is also evidence in rodents that expressing *mHTT,* only during development, and not in later life, still results in neurodegeneration, suggesting that exposing developing neurons to *mHTT* may render them more vulnerable to degeneration as they age.^8^

An important gap in this data has been the availability of single-cell analysis of human fetal brain tissue; in particular of the whole ganglionic eminence (WGE), comprising lateral, medial, and caudal (LGE/MGE/CGE) parts, with the LGE being the main origin of MSNs. Human fetal HD-positive tissue is scarce, and the existence of intact fetal tissue, that allows identification and dissection of WGE, is exceptionally rare with no previous reports of HD-positive human striatal tissues at the time of writing. Here we use single-cell RNA sequencing (scRNAseq) to characterise the gene expression profiles of developing striatal tissue from a human HD positive donor and compare this to an age-matched, non-HD, control fetal sample.

## Materials and Methods

### Detailed methods are included in supplementary materials

**Human Tissue Collection** was conducted with full ethical approval under the Cardiff University HTA licence and Declaration of Helsinki guidelines. Maternal consent was obtained following consent for elective medical termination of pregnancy.

**Dissection** was completed in full for both fetal samples. WGE was processed immediately for sequencing. Remaining WGE and all other tissues were snap-frozen in liquid nitrogen and stored at -80°C. *All brain and peripheral tissues are available for collaborative use. Enquiries to the Brain Repair Group, Cardiff University*.

**CAG-length** was determined using the PanDNA NanoBind kit (Pacific Biosciences), SMRT HiFi amplicon sequencing, Invitrogen Qubit HS dsDNA kit, Pacific Biosciences Sequel IIe.

**Single-cell RNA sequencing** was performed using 10X Genomics Chromium 3’ kits (v3.1) and Illumina NovaSeq6000 (S1, 80,000 reads/cell); processed using Cell Ranger; and analysed in R v4.6 (Seurat v5.5). Cells with <1000 detected genes, unique counts outside 500-6000, or >5% mitochondrial reads were removed. Gene enrichment analysis used adjusted *p*<0.05.

## Results

The HD mutation-carrying (HD) and non-mutation-carrying control (CTRL) fetal samples were retrieved following medical termination of pregnancy. Both were fully intact with no visible disruption of tissue integrity. The samples were male, and well-matched in terms of gestational age, with only 1mm difference in crown-rump-length on direct measurement post-retrieval (Fig. 1A). The cell viabilities for the WGE cell suspensions were 70% and 79% in the HD and CTRL samples respectively.

**Figure 1.**
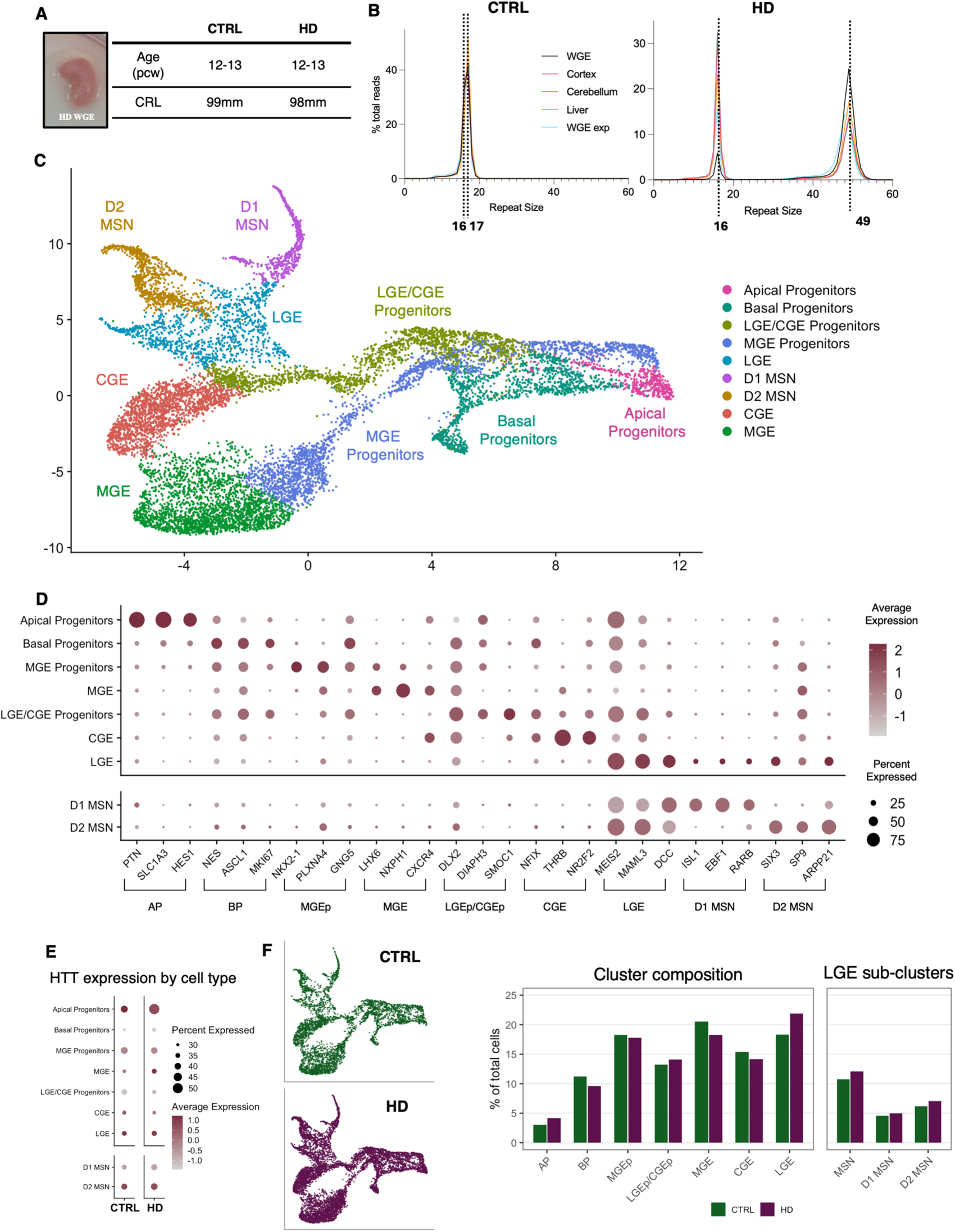
Characterisation of human HD fetal WGE. **(A)** Image of HD WGE at time of dissection and parameters of the fetal tissue samples. The CTRL and HD fetal samples were well-matched for gestational age on both *in utero* ultrasound measurement and direct *ex vivo* crown-rump measurement. CTRL = Control; HD = Huntington’s disease; pcw = post conception weeks; CRL = crown-rump-length. **(B)** Line graphs showing analysis of CAG repeat size in different tissues (WGE, *in vitro* expanded WGE cells, cortex, cerebellum, liver) from CTRL and HD samples. WGE = whole ganglionic eminence; exp = *in vitro* expanded cells. **(C)** UMAP plot showing cluster analysis of single-cell RNA sequencing data from CTRL and HD WGE. Each dot represents a single cell; cells are coloured based on clustering, and annotation is based on gene expression of canonical markers within each cluster. UMAP = Uniform manifold approximation and projection; LGE = lateral ganglionic eminence; CGE = caudal GE; MGE = medial GE; D1 MSN = dopamine receptor type 1 medium spiny projection neuron; D2 MSN = dopamine receptor type 2 MSN. **(D)** Dot plots showing canonical gene expression within the whole population of sequenced cells. The size of the dot represents the percentage of cells within a given cluster expressing a given gene. The colour of the dot represents the average level of expression across the cells expressing the gene. **(E)** Dot plot showing expression of *HTT* in each cluster from both CTRL and HD fetal samples. **(F)** Deconstructed UMAP plots **in C** showing each sample (genotype) individually, and bar charts showing the composition of each annotated cluster in **C** of each sample, as a percentage of total cells; this is displayed on the left for each major cell type in the WGE, and on the right for the key sub-clusters of the LGE (green = CTRL; purple = HD). AP = apical progenitors; BP = basal progenitors. Alt text: Characterisation of human Huntington’s Disease fetal whole ganglionic eminence samples, with subfigures labelled A to F, illustrating tissue dissection images and sample parameters, CAG repeat size analysis across tissue types, single-cell RNA sequencing cluster plots, canonical gene expression dot plots, HTT expression profiles, and deconstructed cell population composition charts comparing control and Huntington’s Disease samples.

### Somatic expansion in the WGE was not detected

The maternal donor was known to have an expanded HD allele (50 (±1) CAG-repeat), and prenatal chorionic villus sampling had revealed the fetus to have inherited the expanded allele. CAG-repeat length was assessed in various brain tissues (WGE, cortex, cerebellum, and 4-week *in vitro* expanded WGE) and liver for both fetal samples. No somatic expansion was seen in any tissue for either allele, in either the HD or CTRL fetal sample (Fig.1B).

### Single-cell RNA sequencing characterisation of HD fetal striatum

We performed scRNAseq on WGE from HD and CTRL fetal samples. Following QC and normalisation, 12,370 viable, good-quality cells clustered into sub-populations of known striatal development. Clustering analysis resulted in nine distinct clusters of well-described striatal cell populations (Fig. 1C): pan-ganglionic eminence (GE) progenitor pool, which sub-divided into apical progenitors (AP) and basal progenitors (BP); more fate-specified progenitor GE cells that branched into MGE progenitors and LGE/CGE progenitors; and more mature cells clustered from their respective progenitor streams into MGE, CGE and LGE, with the LGE subdivision branching further into D1 and D2 MSNs (Fig. 1C). Canonical GE progenitor and fate-committed gene expression analyses showed characteristic, striatal-specific expression patterns of the different GE regions in the cell clusters, illustrated by a dot plot (Fig. 1D) and feature plots (Supplementary Fig. 1). Comparison of expression of *HTT* across the clusters revealed subtle differences between the two samples, with the greatest difference being present in the AP population where a higher proportion of cells expressed *HTT* in the HD sample than the CTRL (Fig. 1E). There were no differences in the proportions of cells in each cluster between genotype (Fig. 1F).

### Differential gene expression analysis revealed specific disturbances across neuro-pathways in HD WGE

There were 2,032 significantly differentially expressed genes (DEGs) between HD and CTRL WGE, with four times more genes upregulated (1,653) than downregulated (379) in HD (Fig. 2A). Visualisation of these DEGs revealed that the majority of the transcriptional profile was similar between genotypes, and that differences were largely driven by subtle perturbations rather than a strong focal defect. Greater magnitude of change in DEGs were more typically observed in genes that were expressed in smaller numbers of cells (Fig. 2B, C).

**Figure 2.**
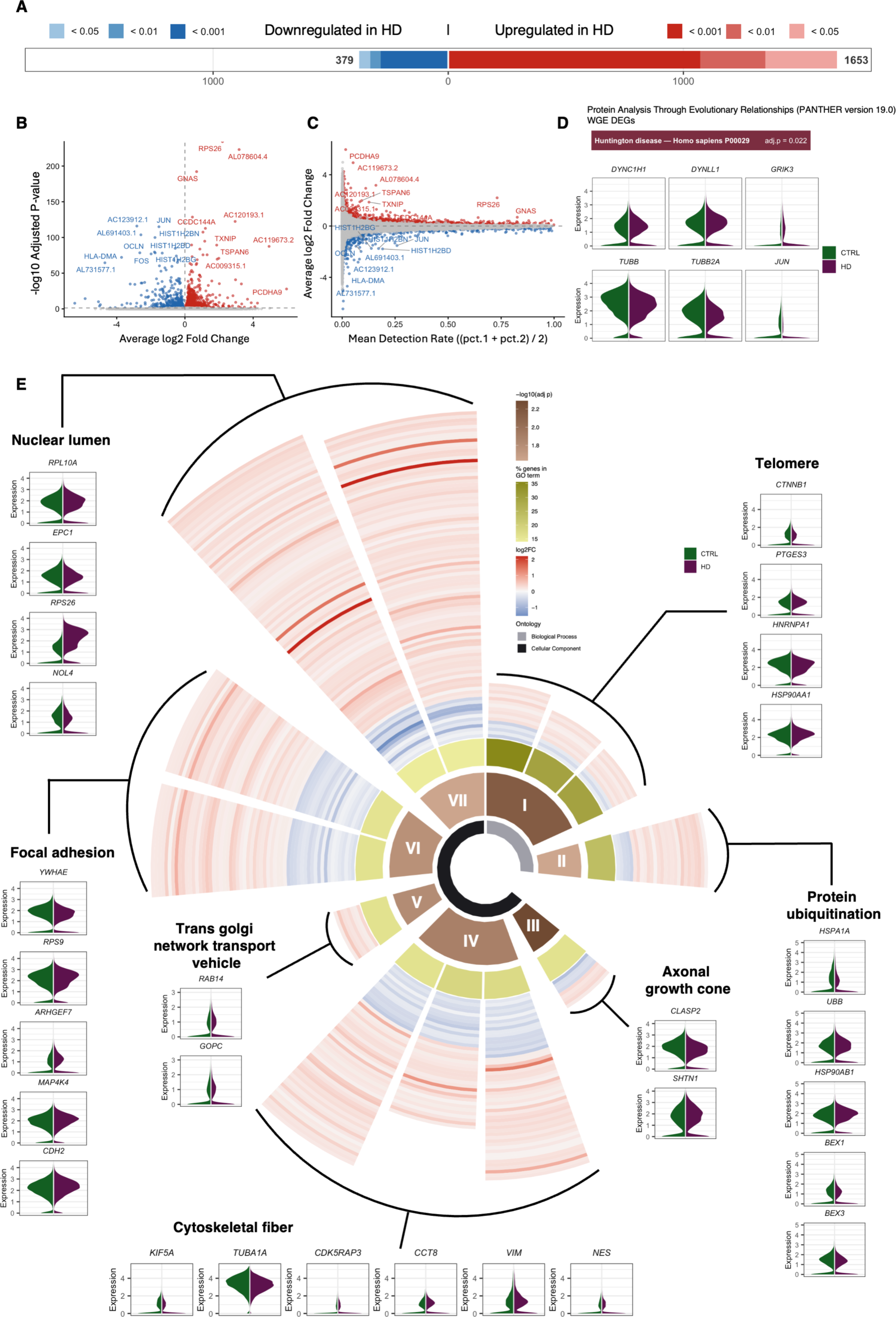
Differentially expressed genes (DEGs) between HD and CTRL WGE. **(A)** Horizontal chart showing the number of DEGs across WGE. Colour gradient indicates threshold of adjusted p (red = upregulated, blue = downregulated; most intense = *p*<0.001, less intense = *p*<0.01, least intense = *p*<0.05). **(B)** Volcano plot showing DEGs across WGE, where fold change (logFC) is plotted against adjusted *p* value (adj *p*) (red = upregulated, blue = downregulated). **(C)** MA plot, showing logFC against the mean detection rate within the population (red = upregulated, blue = downregulated) (MA = minus average), indicating that genes which exhibit greatest fold change were typically seen in cells with the lowest mean detection rates. **(D)** PANTHER (Protein Analysis Through Evolutionary Relationships, version 19.0) pathway analysis yielded “Huntington Disease” as the top significantly enriched term. Corresponding violin plots depicting examples of DEGs from the “Huntington Disease” term (green = CTRL, purple = HD). **(E)** Sunburst plot and corresponding violin plots: The sunburst plot shows gene enrichment analysis (GEA) of all DEGs across the WGE. We explored the GO Biological processes (BioP), Cellular components (CellC) and Molecular function (MolF). In the Sunburst plot: Inner ring, grey = BioP, black = CellC; second ring (I-VII) represents parent categories, defined as the representative term of each semantic similarity group identified by rrvgo, coloured by −log10(adj *<u>p</u>*-value) of all GO terms within that group: I = BioP telomere, II = BioP regulation of protein ubiquitination, III = CellC axonal growth cone, IV = CellC polymeric cytoskeletal fiber, V = CellC Trans-Golgi Network Transport Vesicle, VI = CellC focal adhesion, VII = CellC nuclear lumen; third ring represents the DEGs as a % of the corresponding enriched term gene list: darker = higher %, paler = lower %; the outer-segments represent the log2FC of individual DEGs (red = upregulated, blue = downregulated; intensity represents magnitude). Violin plots showing examples of DEGs underpinning the ontologies shown in the Sunburst plot (green = CTRL; purple = HD). Alt text: Differentially expressed genes between Huntington’s Disease and control whole genome expression samples, with subfigures labelled A to E, illustrating gene expression analyses including a bar chart of DEG counts, volcano and MA plots, pathway enrichment analysis with violin plots, and a sunburst plot with accompanying gene ontology violin plots.

We performed gene enrichment analysis (GEA) on significant DEGs to understand the biological implications of these differences. The only significant term from the Protein Anlaysis Through Evolutionary Relationships (PANTHER) database was “Huntington Disease”, (adjusted *p* value = 0.022), indicating that established disturbances associated with HD are already present in the developing fetal striatum (Fig. 2D). Furthermore, we found disruption of telomere regulation, protein ubiquitination, axonal growth cone, cytoskeletal fiber, trans-golgi network transport vesicle, focal adhesion and nuclear lumen in gene ontology databases (Fig. 2E). The most significant DEG was RPS26 (Fig2. B), with significantly increased expression in the HD WGE compared with CTRL, contributing to disruption of nuclear lumen process. Notably, CDH2 (within focal adhesion) and CTNNB1 (within disrupted telomere, ubiquitination and focal adhesion), were previously shown to be altered in HD-positive developing cortical tissues.^4^ Among the cytoskeletal-associated DEGs were TUBB family members (TUBB, TUBBA1A, TUBB2A, TUBB2B, TUBB3, TUBB4A and TUBB4B); encoding microtubule-related proteins that were consistently downregulated across the HD WGE. Taken together, this data highlights the specific key pathways and cellular functions that are dysregulated within the developing HD striatum.

### Cell type analysis revealed unique patterns of dysregulation within discrete WGE populations

Next, we explored DEGs between HD and CTRL WGE tissues within the cell type subclusters (Fig. 3A). Set intersection analysis of each cell types’ up-/down-regulated DEGs, revealed that HD WGE cell types had unique patterns of gene expression (Fig. 3B). For instance, in the MGE we identified 73 unique genes (55 up- and 18 down-regulated). While some shared gene sets were observed (e.g. LGE and MGE shared 10 up- and 3 down-regulated genes) the majority of the gene sets were not intersectional, suggesting that *mHTT* could be impacting different processes across WGE cell types.

**Figure 3.**
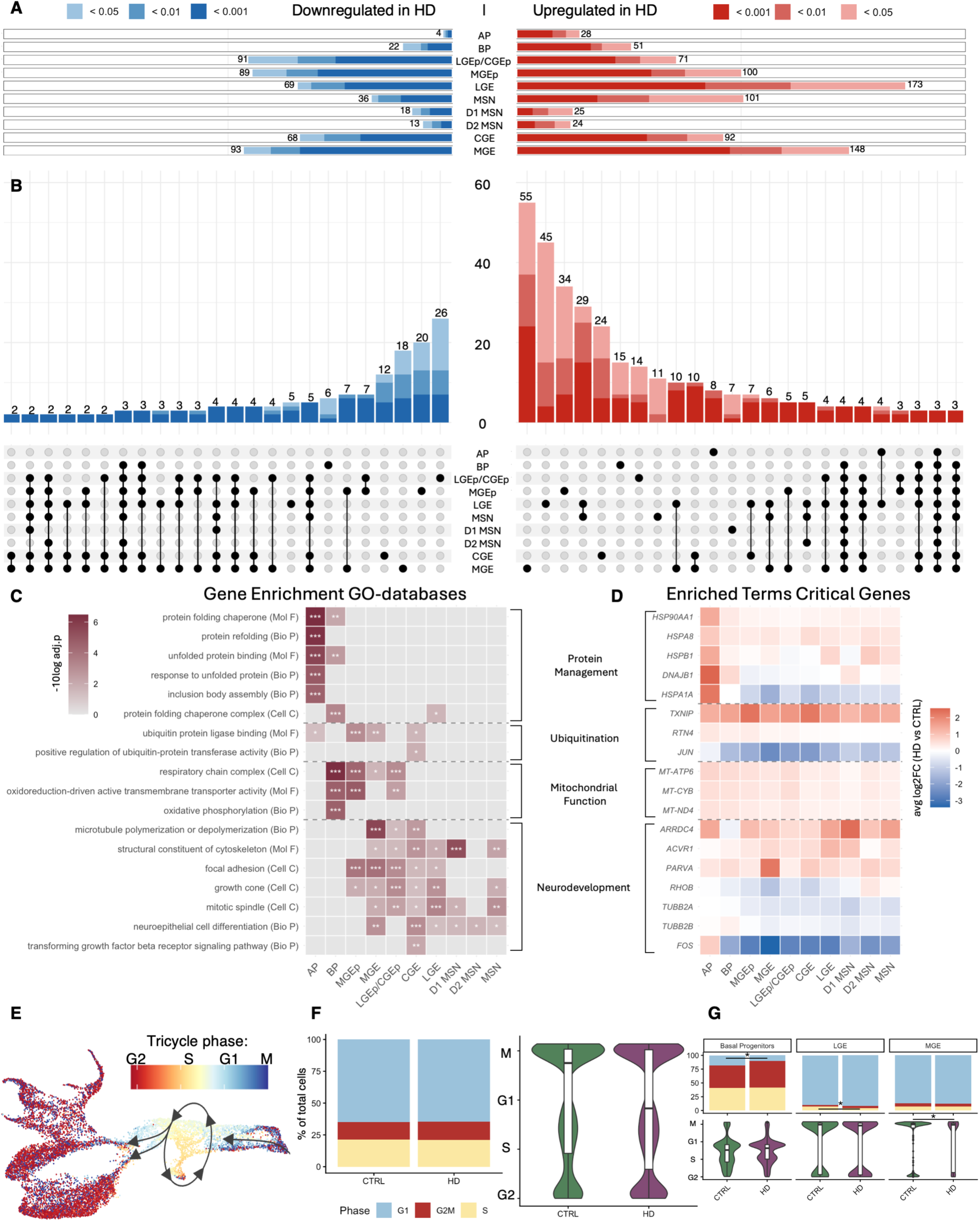
Cell type subcluster analysis between HD and CTRL. **(A)** Horizontal chart showing the number of DEGs in each cluster (from Fig. 1C). Colour gradient indicates threshold of adjusted *p* (red = upregulated, blue = downregulated; most intense = *p*<0.001, less intense = *p*<0.01, least intense = *p*<0.05). AP = Apical progenitors; BP = Basal progenitors; LGEp/CGEp = lateral ganglionic eminence progenitors/caudal ganglionic eminence progenitors; MGEp = medial GE progenitors; MSN = medium spiny projection neuron; D1 MSN = dopamine receptor type 1 MSN; D2 MSN = dopamine receptor type 2 MSN. **(B)** Upset plots showing results of gene set intersection analysis. Gene set intersections are shown in vertical bars with the number of DEGs in that set, positioned over a dot matrix illustrating clusters in which those gene sets are shared. Colour gradient indicates threshold of adjusted *p* (red = upregulated, blue = downregulated; most intense = *p*<0.001, less intense = *p*<0.01, least intense = *p*<0.05). **(C)** Heat map of cell type’s DEG list GEA, illustrating collective contributions to key main terms, *** = *p*<0.001, ** = *p*<0.01, * = *p*<0.05. **(D)** Heat map of average gene expression (log2FC HD vs CTRL) by cell type for critical genes underpinning GEA terms shown in **(C)** (red = upregulated, blue = downregulated; intensity represents magnitude). **(E)** TriCycle analysis of cell cycle phase projected onto UMAP from Fig. 1C. **(F)** Quantification of cell cycle phase across the WGE population expressed as bar chart (Seurat) and violin plot (TriCycle) Showing no change between CTRL and HD. **(G)** Quantification of cell cycle phase within cell types, expressed as bar chart (Seurat) and violin plot (TriCycle) (green = CTRL; purple = HD), * = *p*<0.05. Alt text: Cell type subcluster differentially expressed gene analysis between Huntington’s Disease and control whole ganglionic eminence samples, with subfigures labelled A to G, illustrating cluster-specific differential gene expression bar charts, gene set intersection upset plots, gene enrichment analysis heatmaps, average gene expression heatmaps by cell type, cell cycle phase projections onto dimensionality reduction plots, and cell cycle phase quantification across whole population and individual cell types comparing control and Huntington’s Disease samples.

To better understand the impact of these cell type specific variations, we conducted GEA on each cell types’ DEG list. We identified over 100 significantly enriched terms across these cell types (Supplementary Fig. 2), the majority of which were related to three major themes: protein regulation including folding/refolding, and ubiquitination; mitochondrial function of oxidation and ATP synthesis; and neuronal development regulation, organisation and migration (Supplementary Fig. 3). We observed a clear trend for genes associated with protein regulation to exhibit heightened expression in AP populations of the HD tissue (Fig. 3C, D), which notably coincided with broader *HTT* expression compared to CTRL within this cell type (Fig. 1E). Greater expression of genes regulating mitochondrial function was observed across all HD populations, but particularly in the wider progenitor pool (Fig. 3C, D). Finally, DEGs associated with neuronal development were significantly dysregulated in maturing tissues beyond the apical/basal fate, where neuronal fate commitment is being established (Fig. 3C, D). Collectively these results demonstrate consistent and specific alterations in subpopulations of the HD WGE.

### Analysis of cell-cycling between HD and CTRL WGE tissues

Alterations to cell-cycling have previously been identified in HD development.^4,9^ Subsequently, we specifically explored the cell-cycle process across our WGE tissues (Fig. 3E-G). Across the WGE, we observed active mitosis, with a major cycling stream occurring within the earlier BP population (Fig. 3E). While, no differences in cell numbers within different phases were observed across the global population (Fig. 3F), we did identify significant differences of cell-cycle phases within the large cycling BP population, and both the LGE and MGE committed cells, suggesting alterations to cell-cycling behaviours in these discrete cell types (Fig. 3G).

### Do disease specific HD signatures already exist in the developing HD striatum?

To investigate whether our findings in the HD WGE reflect changes seen in the adult HD striatum (comprised of caudate (Cu), putamen and nucleus accumbens (nA)), we compared our DEGs to publicly available postmortem (PM) datasets of these brain regions^10^. We focused on Cu tissue Vonsattel^11^ Grades 0-1 (Hodges et al.^12^); and Cu and nA tissue Vonsattel Grades 2-4 (Paryani et al.^13^). We found 3,651 DEGs in the HD Cu grades 0-1; 2,112 in the HD Cu grades 2-4; and 666 in the HD nA. We aligned DEGs from WGE and PM tissue to identify shared DEGs: the largest overlap was with Cu grades 2-4 (398 genes), followed by Cu grades 0-1 (285 genes) and nA (62 genes) (Fig. 4A). Neuron projection, axon guidance, vesicles, and nuclear speck were identified as some of the altered processes from the shared DEG-list between HD WGE and adult PM tissue (Supplementary Fig. 4).

**Figure 4.**
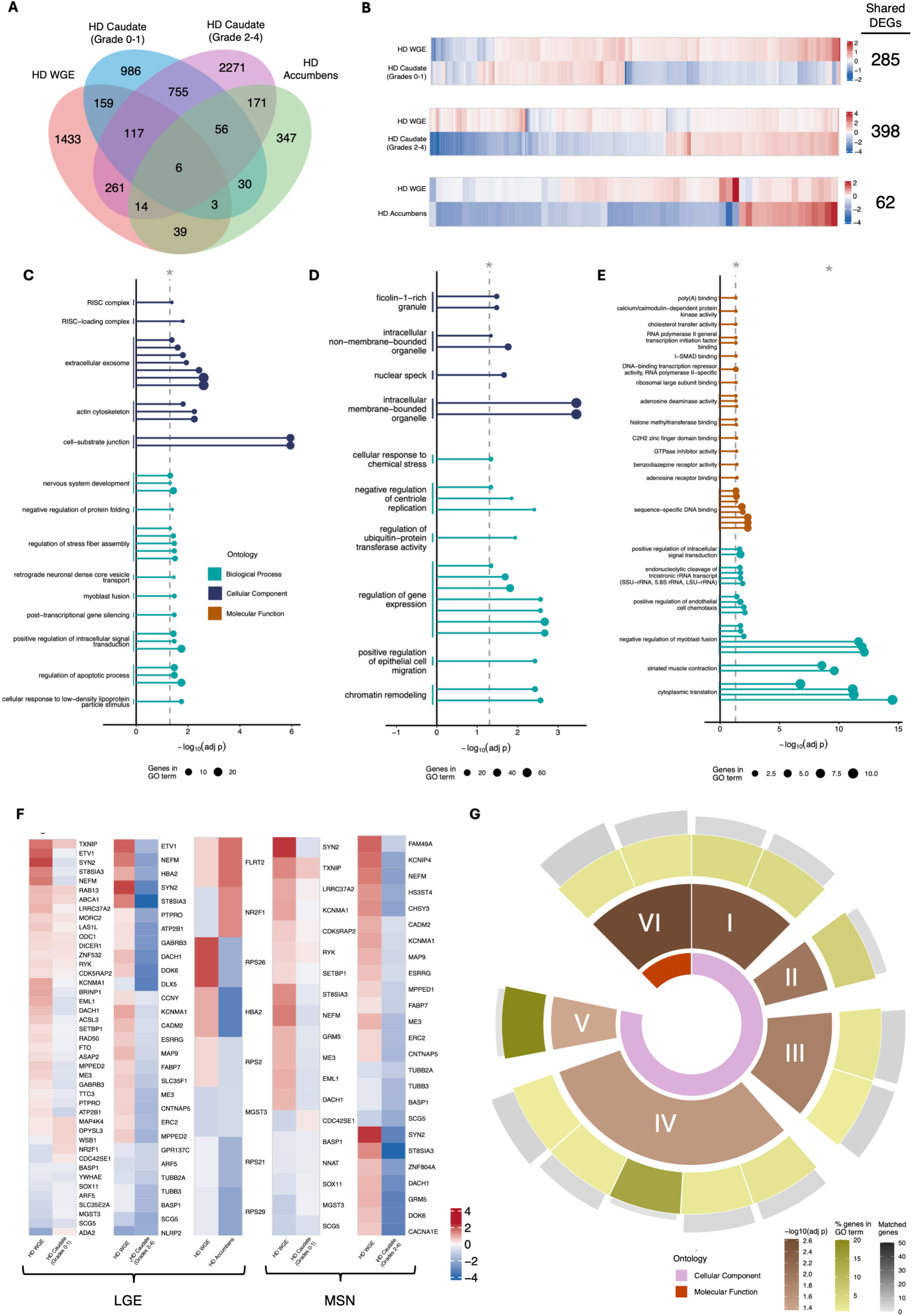
Comparison of DEGs between HD WGE and adult HD postmortem tissue. **(A)** Venn diagram showing the overlap of DEGs identified in HD WGE, with adult HD postmortem (PM) caudate (Cu) Vonstattel grades 0–1, Cu grades 2–4, and nucleus accumbens (nA) grades 2-4. DEGs within a dataset were identified independently by comparison to region- and age-matched controls. **(B)** Heatmaps illustrating the LogFC direction of shared DEGs between HD WGE and adult PM HD Cu grades 0-1, Cu grades 2-4 and nA grades 2-4 (red = upregulated, blue = downregulated; intensity represents magnitude). **(C), (D), (E)** Lollipop plots of GEA of the shared DEGs between WGE and adult HD PM tissue for ontologies BioP, MolF and CellC. GEA on overlapping DEGs that share the same direction in WGE and **(C)** adult Cu grades 0-1; **(D)** adult Cu grades 2-4; and **(E)** adult nA grades 2-4. The representative ontology term of each group is shown on the y axis. Each lollipop represents an ontology term: length represents −log10(adj *p*-value) of that term and size of the head represents the number of genes. Vertical dashed line represents *p*=0.05. **(F)** Heatmaps showing shared DEGs from the LGE (left hand side) and MSN (right hand side) clusters with adult Cu grades 0-1, adult Cu grades 2-4 and adult nA grades 2-4 (red = upregulated, blue = downregulated; intensity represents magnitude). **(G)** Sunburst plot showing GEA of shared DEGs between the WGE LGE cluster and the adult HD Cu grades 2-4. Inner ring: orange = MolF, red = CellC; second ring (I-VI) represents parent categories, defined as the representative term of each group, coloured by −log10(adj *<u>p</u>*-value) of all GO terms within that group; I = synaptic vesicle membrane, II = GABA-ergic synapse, III = axon, IV = mitotic spindle, V = neurofibrillary tangle, VI = guanyl ribonucleotide binding; third ring represents the DEGs as a % of the corresponding enriched term gene list (darker = higher %, paler = lower %); outer segments represent number of genes within an ontology term shown by length and colour. Alt text: Comparison of differentially expressed genes between Huntington’s Disease whole ganglionic eminence and adult Huntington’s Disease postmortem tissue across multiple brain regions and disease grades, with subfigures labelled A to G, illustrating overlapping gene sets via Venn diagram, fold change direction heatmaps, gene enrichment lollipop plots across biological process, molecular function and cellular component ontologies for caudate and nucleus accumbens tissue at varying disease grades, cell-type-specific shared gene expression heatmaps, and a sunburst plot of gene enrichment analysis for lateral ganglionic eminence cluster genes shared with adult caudate tissue.

Directional analysis of the shared DEGs revealed that 124 (Cu grades 0-1), 175 (Cu grades 2-4) and 33 (nA) DEGs remained consistent in their direction of change between the HD WGE and adult HD brain (Fig. 4B). GEA of these shared-direction genes identified pathways of protein folding and management, gene expression and stress response (Fig. 4C-E). Additionally, pathways enriched for changed-direction-DEGs were explored. For instance, DEGs downregulated in WGE and upregulated in PM Cu grades 2-4 yielded ontology terms associated with dysregulated mitochondrial function, regulation of interleukin production, cell regulation and growth transition (Supplementary Fig. 5).

Next, we explored shared DEGs between WGE cell types (LGE, MSN, CGE clusters) and the HD PM tissue. We identified 43 (Cu grades 0-1) 29 (Cu grades 2-4) and 8 (nA) DEGs shared with the LGE cell type; 19 (Cu grades 0-1) and 25 (Cu grades 2-4) DEGs with the MSN cell type; and 6 DEGs between the CGE cell type and the nA (Fig. 4F, Supplementary Fig. 6). GEA revealed cell type specific alterations, for example DEGs from the Cu (grades 2-4) shared with the LGE were associated with GABA-ergic synapse, axon, mitotic spindle, neurofibrillary tangle, guanyl ribonucleotide binding and synaptic vesicle membrane (Fig. 4G, additional clusters in Supplementary Fig. 7). Overall, these data demonstrate that some disturbances in gene expression observed in adult HD aren already present in the developing brain and are probably maintained across the disease lifespan.

## Discussion

We present the first scRNAseq analysis of the developing human HD fetal striatum (WGE), offering novel insight into the earliest effects of the disease on the most impacted cells. A key limitation is the use of a single HD-positive fetal sample; obtaining this exceptionally rare sample required ten years of academic resource and consistent engagement with clinical partners. Some mitigation was achieved through close matching to a control sample (i.e. sex-matched, age-matched with almost identical CRLs, and the healthy allele being well-matched). To minimise non-biological variation between samples, all technical aspects were conducted under identical conditions, and were processed for sequencing together. Importantly, many of our findings are corroborated by evidence from other tissues and disease-models.

We first assessed somatic expansion of the *mHTT* CAG-repeat. This is present years before clinical onset, but the time of its initiation remains unknown. Kacher et al.^14^ found no somatic expansion in human fetal HD cortex, and here we demonstrate absence of somatic expansion in HD WGE, other fetal brain tissues, and liver; extending previous findings and confirming that somatic expansion has not begun at this early developmental stage.

ScRNAseq revealed nearly four times more upregulated than downregulated DEGs in the HD WGE. Most changes were either subtle across many cells, or pronounced across few cells, with only a handful of widely-expressed genes exhibiting large differences between the genotypes. Cluster analysis revealed cell type-specific transcriptional dysregulation, frequently related to known dysregulated pathways in HD. For example, increased expression of heat-shock protein genes (relating to protein folding and management) occurred only in HD AP, and coincided with more widespread *HTT* expression in this cluster compared to CTRL. Heat-shock proteins are known to be disrupted in HD,^15,16^ and such cluster-specific transcriptional changes may provide clues to the cell-specific nature of HD pathology.

Some disruptions uncovered in the HD WGE correspond to deficits in development preciously found using different systems. For example, cell-cycle perturbations were reported in human fetal cortex^4^ and mouse models^9^; and axonal growth, as an altered ontology, aligns with HD mouse data demonstrating disorganised cortical axonal growth cones.^17^ Transcriptional dysregulation patterns also substantially aligned with those reported in adult HD tissues. For example, telomere length is shorter in HD patients^18^ and strongly correlates with estimated time to clinical onset,^19^ and the critical telomere-associated gene, CTNNB1 is both downregulated in our study and disrupted in HD fetal cortex.^4^ However, not all transcriptional profiles were mirrored in adult human PM tissue and some were reversed, likely reflecting ageing processes, compensatory mechanisms, and technical limitations inherent to such comparisons.

We believe the data presented here, alongside additional tissues still available for analysis, are an important resource to begin understanding whether, and how abnormalities present in the fetus impact the degenerative process in HD. There is an imperative to share such rare samples and to continue, and extend, efforts to collect further samples. Understanding early brain development in HD is important for many reasons, for example in the identification of biomarkers, and is critical to guiding timeframes and potential targets for therapeutic interventions aimed at delaying or halting progression of the disease.

## Data Availability

The single-cell RNA sequencing data generated in this study have been deposited in ArrayExpress (E-MTAB-17220) and will be made publicly available upon publication.

A complete list of the DEGs and enriched terms can be found in Supplementary Tables 1-12.

## Acknowledgements

We are especially grateful to research midwife Jan Phipps, for working with us and coordinating things at such short notice. Thanks to Abigail Taylor, Chris Tattersall, Amie Jordan, Helen Falconer and their R&D teams; members of the SWIFT team from University Hospital Wales, Cardiff; ANTC, Cardiff. We thank the donors for the donation of fetal material to our research study.

## Abbreviations

Age pcw: gestational age in weeks estimated post conception
AP: Apical progenitors
BP: Basal progenitors
CGE: caudal ganglionic eminence
CRL: crown-rump-length
CTRL: indicates the fetal sample carrying two non-expanded alleles
Cu: caudate
D1: dopamine receptor type 1
D2: dopamine receptor type 2
DEGs: differentially expressed genes
GEA: gene enrichment analysis
HD: Huntington’s disease; indicates the sample carrying the expanded huntingtin allele
LGE: lateral ganglionic eminence
MGE: medial ganglionic eminence
mHTT: mutant Huntingtin
MSN: medium spiny projection neuron
nA: nucleus accumbens
PM: postmortem
scRNAseq: single-cell RNA sequencing
UMAP: Uniform manifold approximation and projection
WGE: whole ganglionic eminence

## Supplementary material

Supplementary material is available at *Brain* online.

Supplementary Table 1. Global significant DEGs

Supplementary Table 2. Apical progenitor DEGs

Supplementary Table 3. Basal progenitor DEGs

Supplementary Table 4. MGE progenitor DEGs

Supplementary Table 5. LGE progenitor-CGE progenitor DEGs

Supplementary Table 6. LGE DEGs

Supplementary Table 7. MGE DEGs

Supplementary Table 8. MSN DEGs

Supplementary Table 9. D1 MSN DEGs

Supplementary Table 10. D2 MSN DEGs

Supplementary Table 11. Enriched Biological processes from Global sig DEGs

Supplementary Table 12. Enriched Cellular Components from Global sig DEGs

Supplementary Fig 1. Feature Plots depicting canonical gene expression across WGE populations

Supplementary Fig 2. Heatmap of GEA results from cluster analysis

Supplementary Fig 3. Treemap of enriched terms identified in cell type cluster GEA.

Supplementary Fig 4. Sunburst plots from GEA on shared DEGs between HD WGE and HD PM tissues

Supplementary Fig 5. Lollipop plots of GEA changed direction DEGs between HD WGE and HD PM tissues

Supplementary Fig 6. Heatmap of shared DEGs between HD WGE CGE cell types and nA PM

Supplementary Fig 7. Sunburst plots from GEA on shared DEGs between HD LGE, MSN, and CGE with HD PM Cu and nA tissues

## References

1. Bates GP, Dorsey R, Gusella JF, et al. Huntington disease. Nat Rev Dis Primers. 2015;1:15005.

2. Scahill RI, Zeun P, Osborne-Crowley K, et al. Biological and clinical characteristics of gene carriers far from predicted onset in the Huntington’s disease Young Adult Study (HD-YAS): a cross-sectional analysis. Lancet Neurol. 2020;19(6):502–512.

3. Ciosi M, Maxwell A, Cumming SA, et al. A genetic association study of glutamine-encoding DNA sequence structures, somatic CAG expansion, and DNA repair gene variants, with Huntington disease clinical outcomes. EBioMedicine. 2019;48:568–580.

4. Barnat M, Capizzi M, Aparicio E, et al. Huntington’s disease alters human neurodevelopment. Science. 2020;369(6505):787–793.

5. Humbert S, Barnat M. Huntington’s disease and brain development. C R Biol. 2022;345(2):77–90.

6. Ratié L, Humbert S. A developmental component to Huntington’s disease. Rev Neurol. 2024;180(5):357–362.

7. Ruzo A, Croft GF, Metzger JJ, et al. Chromosomal instability during neurogenesis in Huntington’s disease. Development. 2018;145(2):156844.

8. Molero AE, Arteaga-Bracho EE, Chen CH, et al. Selective expression of mutant huntingtin during development recapitulates characteristic features of Huntington’s disease. Proc Natl Acad of Sci U S A. 2016;113(20):5736–5741.

9. Molero AE, Gokhan S, Gonzalez S, Molero AE, Gokhan S, Gonzalez S. Impairment of developmental stem cell-mediated striatal neurogenesis and pluripotency genes in a knock-in model of Huntington’s disease. Proc Natl Acad of Sci U S A. 2009;106(51):21900–21905.

10. Wichterle H, Turnbull DH, Nery S, Wichterle H, Turnbull DH, Nery S. In utero fate mapping reveals distinct migratory pathways and fates of neurons born in the mammalian basal forebrain. Development. 2001;128(19):3759–3771.

11. Vonsattel JP, Myers RH, Stevens TJ, Ferrante RJ, Bird ED, Richardson Jr EP. Neuropathological classification of Huntington’s disease. J Neuropathol Exp Neurol. 1985;44(6):559–577.

12. Hodges A, Strand AD, Aragaki AK, et al. Regional and cellular gene expression changes in human Huntington’s disease brain. Hum Mol Genet. 2006;15(6):965–977.

13. Paryani F, Kwon JS, Ng CW, et al. Multi-omic analysis of Huntington’s disease reveals a compensatory astrocyte state. Nat Commun. 2024;15(1):6742.

14. Kacher R, Lejeune FX, Noel S, et al. Propensity for somatic expansion increases over the course of life in Huntington disease. elife. 2021;10:e64674.

15. Chafekar SM, Duennwald ML. Impaired heat shock response in cells expressing full-length polyglutamine-expanded huntingtin. PloS One. 2012;7(5):e37929.

16. Gomez-Pastor R, Burchfiel ET, Neef DW, et al. Abnormal degradation of the neuronal stress-protective transcription factor HSF1 in Huntington’s disease. Nat Commun. 2017;8(1):14405.

17. Capizzi M, Carpentier R, Denarier E, et al. Developmental defects in Huntington’s disease shos that axonal growth and microtubule reorganization require NUMA1. Neuron. 2022;110(1):36–50.

18. Kota LN, Bharath S, Purushottam M, et al. Reduced telomere length in neurodegenerative disorders may suggest shared biology. J Neuropsychiatry Clin Neurosci. 2015;27(2):e92–96.

19. Scarabino D, Veneziano L, Mantuano E, et al. Leukocyte telomere length as potential biomarker of HD progression: a follow-up study. Int J Mol Sci. 2022;23(21):13449.

