## Supplementary Methods for "Cell-specific transcription dysregulation in human Huntington’s disease-positive developing striatum"

**Detailed Materials and Methods**

**Human Tissue Collection: Ethics and Approvals**: This study was conducted with full ethical approval (HD positive fetal tissue was collected under project 13-WA-0361; control fetal sample was obtained via SWIFT-RTB 23/WA/0115), in compliance with HTA regulations under the Cardiff University HTA licence (licence number 12422 ) and under guidelines set out in the Declaration of Helsinki. Consent for tissue donation to the research study was obtained following the consent of the maternal donor to undergo the elective medical termination of pregnancy.

**Dissection:** Following passing, the tissue was transferred to a pot with Hibernate E medium and stored at 4°C. Dissection was performed in Hibernate E medium, with a full dissection of each fetus. One whole ganglionic eminence (WGE), the primordial striatum, from each sample was used to generate a single-cell suspension. Unused cell suspension was cryopreserved and all other dissected tissues were stored at -80°C, for future use. All other dissected tissue pieces were either stored in Hibernate E medium at 4°C or were snap-frozen in liquid nitrogen and stored at -80°C.

**Cell preparation**: Dissected WGE tissue was dissociated using TrypLE Express with Dornase alfa (20 units dornase alfa per ml TrypLE Express). Following incubation at 37°C for 10 minutes the TrypLE+Dornase alfa solution was removed and the tissue was washed three times in DMEM+Dornase alfa (20 units dornase alfa per ml DMEM). In the third wash, the tissue was manually triturated by pipetting 10-20 times to generate a single-cell suspension. Cells were counted and viability assessed using Trypan blue. The cell viabilities for the WGE cell suspensions were 70% (HD sample) and 79% (CTRL sample).

**CAG repeat:** Prior to discussing informed consent for donation to this project, the potential HD pregnancy was known to be carrying the expanded CAG repeat via a diagnostic, prenatal chorionic villus sampling test. The maternal donor of the HD positive tissue was a carrier of the CAG expansion with a repeat length of 50. The CAG repeat length on the paternal side for the HD positive tissue sample and for both the maternal and paternal lineages for the CTRL tissue sample used in this study, were unknown. We sought to determine the CAG repeat length in tissues from both fetal samples performing genome sequencing on tissue pieces from each donor (frozen, stored tissue samples from cortex, cerebellum and liver, cryo-preserved cells from WGE cell suspension, and fixed cells from *in vitro* expanded WGE cells). For CTRL sample, the CAG size was 17 in WGE, *in vitro* expanded WGE, cerebellum and liver, and 16 in cortex. For the HD sample, there were 16 CAG repeats on the unexpanded allele for all tissues, and on the expanded allele, there were 49 CAG repeats in the WGE, cortex, cerebellum and liver tissues and 48 CAG repeats in the *in vitro* expanded WGE. This analysis revealed no evidence of expansion of the CAG repeat within these tissues at this age.

Frozen foetal tissue was pulverised, and genomic DNA was extracted with the PanDNA NanoBind kit (Pacific Biosciences). Repeats at the HTT locus were sequenced using SMRT HiFi amplicon sequencing. Three PCRs (3 x 50 µl volume) per sample were combined and cleaned with AMPure PB beads according to PacBio’s barcoded overhang adapter protocol (Pacific Biosciences #101-791-700). The amplicon library preparation was done according to the manufacturer’s instructions (Pacific Biosciences). Library concentration was quantified using the Invitrogen Qubit HS dsDNA kit. Sample pools were sequenced using the Pacific Biosciences Sequel IIe.

**Single-cell RNA sequencing**: Single-cell RNA sequencing was performed using the 10X Genomics platform on single-cell suspensions of CTRL and HD WGE tissue (10,000 cells per sample at 1000 cells per μl) according to Manufacturers’ instructions (Chromium Next GEM Single Cell 3’ Reagent Kits, 10X Genomics). Cells were sequenced using the Illumina NovaSeq 6000, at 80,000 reads per cell, with paired-end sequencing, 28bp Read1 / 90bp Read2 / 10bp+10bp dual index reads, S1 flow cell mode. Raw data files (Fastq) were demultiplexed, processed, aligned (refdata-gex-GRCh38-2020-A) and quantified using Cell Ranger Single-Cell Software Suite (10x Genomics).

**Data Processing:** Downstream analysis was conducted using Seurat version 5.5 in R version 4.6, with additional packages: clusterProfiler_4.20.0, org.Hs.eg.db_3.23.1, GOSemSim_2.38.0, TriCycle 1.2, Rrvgo 1.24, ComplexUpset_1.3.3, tibble_3.3.1, tidyr_1.3.2, stringr_1.6.0, dplyr_1.2.1,and EVenn. Quality control, data normalisation, and dimensionality reduction were conducted using standard Seurat workflows and packages. Cells with less than 1000 detected genes and with unique counts outside the range of 500-6000 were excluded. Any cells where the mitochondrial count exceeded 5% were removed. Cells were clustered using Seurat's graph-based Louvain clustering algorithm. To determine an appropriate clustering resolution, clustering was performed across a range of resolutions (0.01–1.2), and the resulting cluster solutions were evaluated against the expression of canonical marker genes for ganglionic eminence-derived progenitor and neuronal populations (See Figure 1D and Supplementary Figure 1). Cluster identities were assigned based on concordance between marker gene expression and the cell-type definitions reported in Shi et al., 2021.

**Differentially expressed gene analysis:** DEGs between HD and control conditions were identified using Seurat, applying the Wilcoxon rank-sum test. Comparisons were performed both globally (across the entire dataset) and within each individual cell type cluster, with HD set as the comparison group relative to control. To allow for unfiltered downstream analysis, no minimum percentage expression or log fold-change threshold was applied at the testing stage; all detected genes were therefore tested regardless of expression magnitude or prevalence. Resulting p-values were adjusted for multiple comparisons using the Bonferroni correction, and genes with an adjusted p-values below 0.05 were considered statistically significant. For the combined D1 and D2 medium spiny neuron (MSN) population, cells from both subtypes were grouped prior to differential expression testing to increase statistical power.

**Gene enrichment analysis:** Gene enrichment analysis was performed on significant DEGs (up- and down-regulated genes were combined) identified for each cluster, using Enrichr. Enrichment was assessed independently across all three GO domains: Biological Process, Molecular Function, and Cellular Component. P-values were adjusted for multiple comparisons using the Benjamini-Hochberg procedure, and enriched terms with an adjusted p-value below 0.05 were considered statistically significant. To reduce redundancy arising from semantically overlapping GO terms, significant terms within each cluster and GO domain were further refined.

**Postmortem dataset analysis:** Data for the caudate (grade 2-4) and nucleus accumbens (grade 2-4) was taken from Paryani et al., 2024 (GSE242195) and data for caudate (grade 0-1) originated from Hodges et al., 2006 (GSE3790).

**Bulk-RNA analysis:** Gene expression data for the caudate (grade 2–4) and nucleus accumbens (grade 2–4) comparisons were generated using bulk RNA sequencing. The original dataset contained juvenile HD samples, for our analysis did not include these samples as their CAG repeat did not align with our fetal sample. There were 6 controls and 6 HD samples for the caudate comparison and 4 controls and 6 HD samples for the nucleus accumbens comparison. These regions were graded as Vonstattel grades 2, 3 or 4. DEGs were determined from read counts using DESeq2. Prior to differential expression testing, lowly expressed genes were filtered, retaining only genes with a summed count of at least 10 across all samples. Sex was included as a covariate in the DESeq2 design formula to account for sex-related variation in gene expression. Genes were considered differentially expressed if they had an adjusted p-value of <0.05 (Benjamini-Hochberg).

**Microarray analysis:** Gene expression data for the caudate (grade 0-1) comparison was generated using bulk RNA sequencing. The original analysis by Hodges et al., 2006 contained PM tissue from grades 0,1,2,3 and 4. Prior to differential expression analysis, lowly expressed probes were filtered, retaining only probes with an expression level above 100 in at least 3 samples. Differential expression analysis was performed using limma, with a linear model fit followed by empirical Bayes moderation. Genes were considered differentially expressed at an adjusted p-value of <0.05 (Benjamini-Hochberg).
