## Supplementary Figures for "Cell-specific transcription dysregulation in human Huntington’s disease-positive developing striatum"

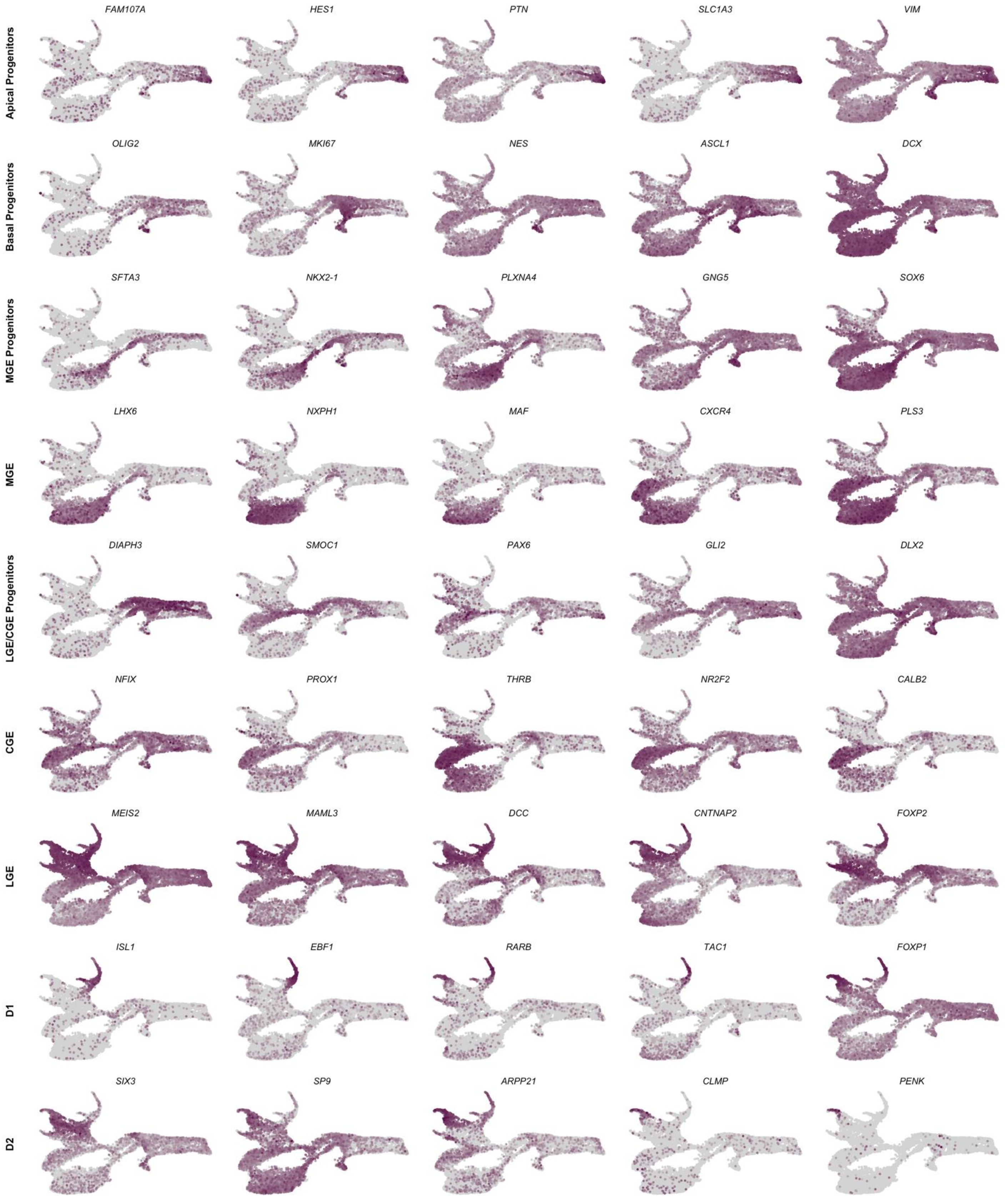


Supplementary Figure 1.
Feature Plots depicting canonical gene expression across WGE populations

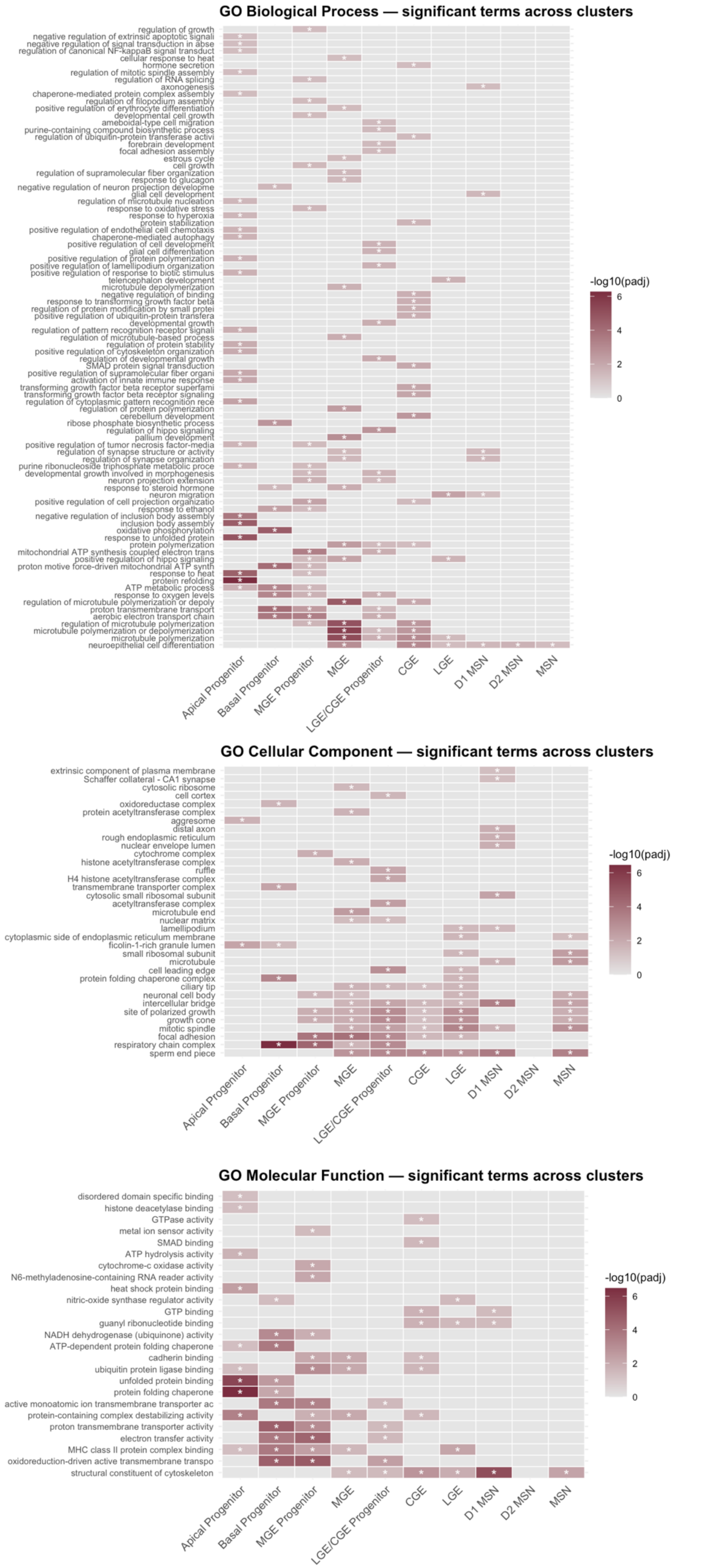


Supplementary Figure 2
Heatmap of GEA results from cluster analysis

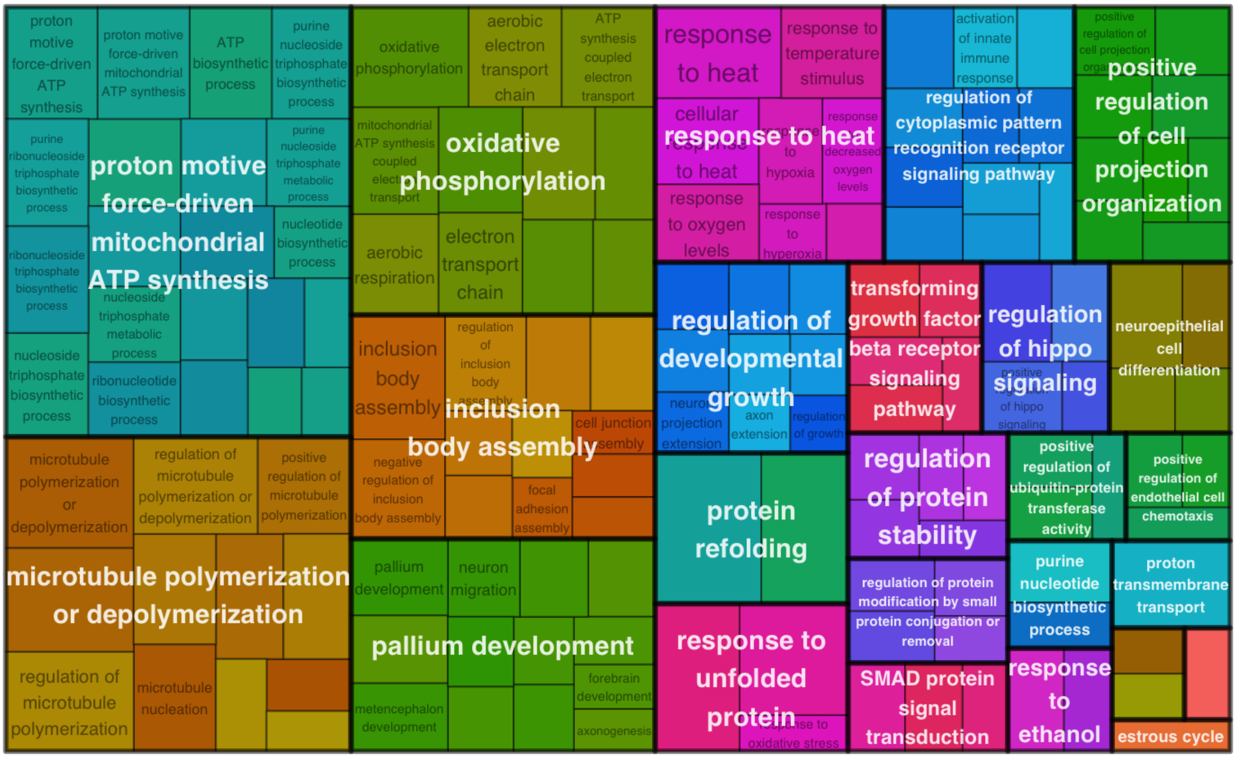


Supplementary Figure 3
Treemap of enriched terms identified in cell type cluster GEA.

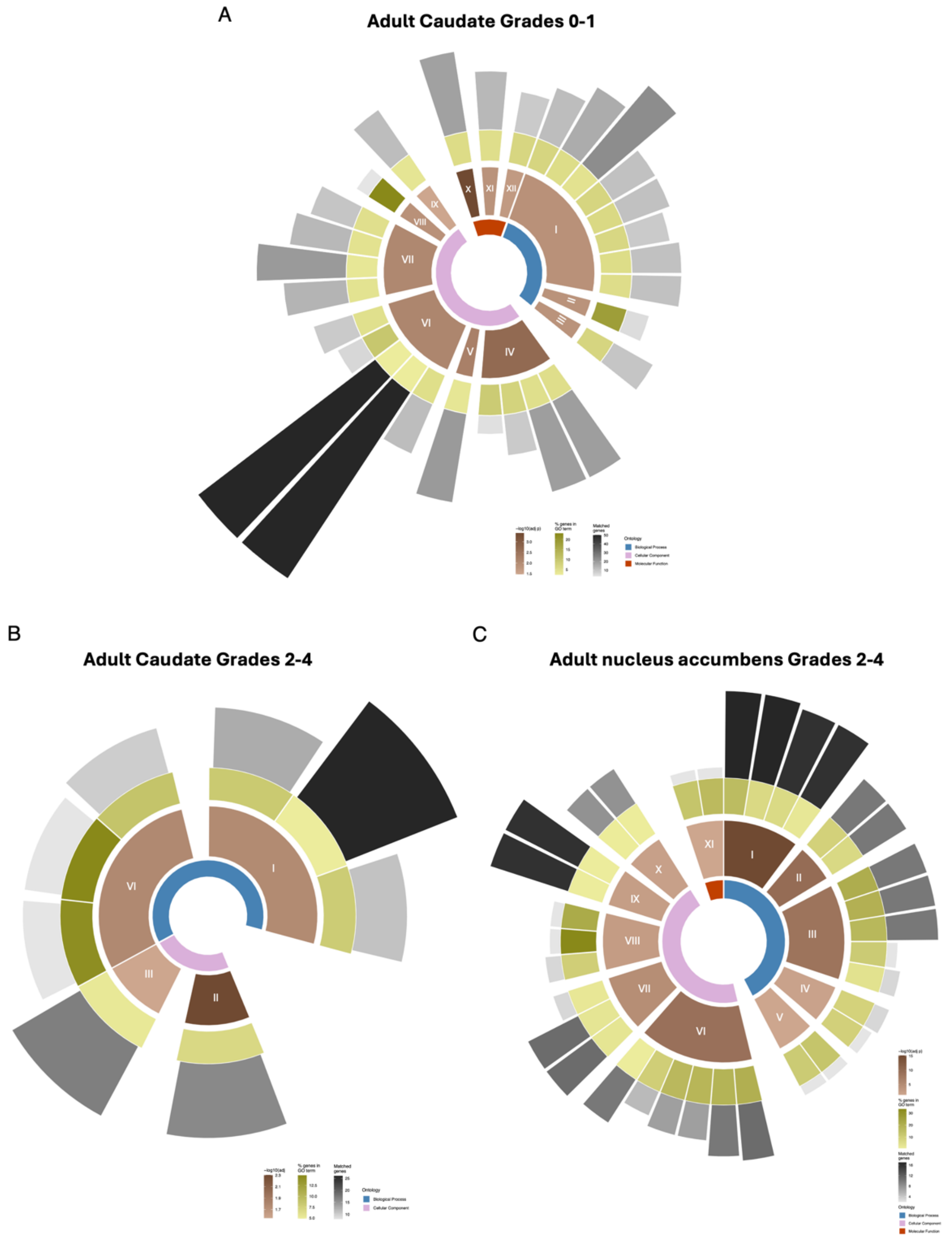


**Supplementary Figure 4: Sunburst plots from GEA on shared DEGs between HD WGE and HD PM tissues.** Sunburst plot showing GEA of shared DEGs between the WGE LGE cluster and the adult HD caudate grades 2-4. Inner ring: blue = BioP orange = MolF, pink = CellC; second ring (I-VI) represents parent categories, defined as the representative term of each group, coloured by −log10(adj p-value) of all GO terms within that group; third ring represents the DEGs as a % of the corresponding enriched term gene list (darker = higher %, paler = lower %); outer segments represent number of genes within an ontology term shown by length and colour. **A s**hared DEGs between adult Cu grades 0-1: I = brain development, II = cellular response to low-density lipoprotein particle stimulus, III = modulation of chemical synaptic transmission, IIII = focal adhesion, IV = neuron projection, V = vesicle, VI = actin cytoskeleton, VII = beta-catenin-TCF complex, VIII = nuclear speck, IX = microtubule binding, X= ion channel regulator activity. **B** shared DEGs between adult Cu grades 2-4: I =negative regulation of protein modulation by small protein conjugation or removal, II = axon guidance, III = vesicle, IIII = nuclear speck. **C** shared DEGs between adult nA grades 2-4: I = cytoplasmic translation, II = striated muscle contraction, III = negtaive regulation of myoblast fusion, IIII = negative regulation of RNA splicing, IV = positive regulation of endothelial cell chemotaxis, V = large ribosomal subunit, VI = focal adhesion, VII = cytoplasmic side of endoplasmic reticulum membane, VIII = extracellular exosome, IX =nucleolus, X= ubiquitin-protein transferase inhibitor activity.

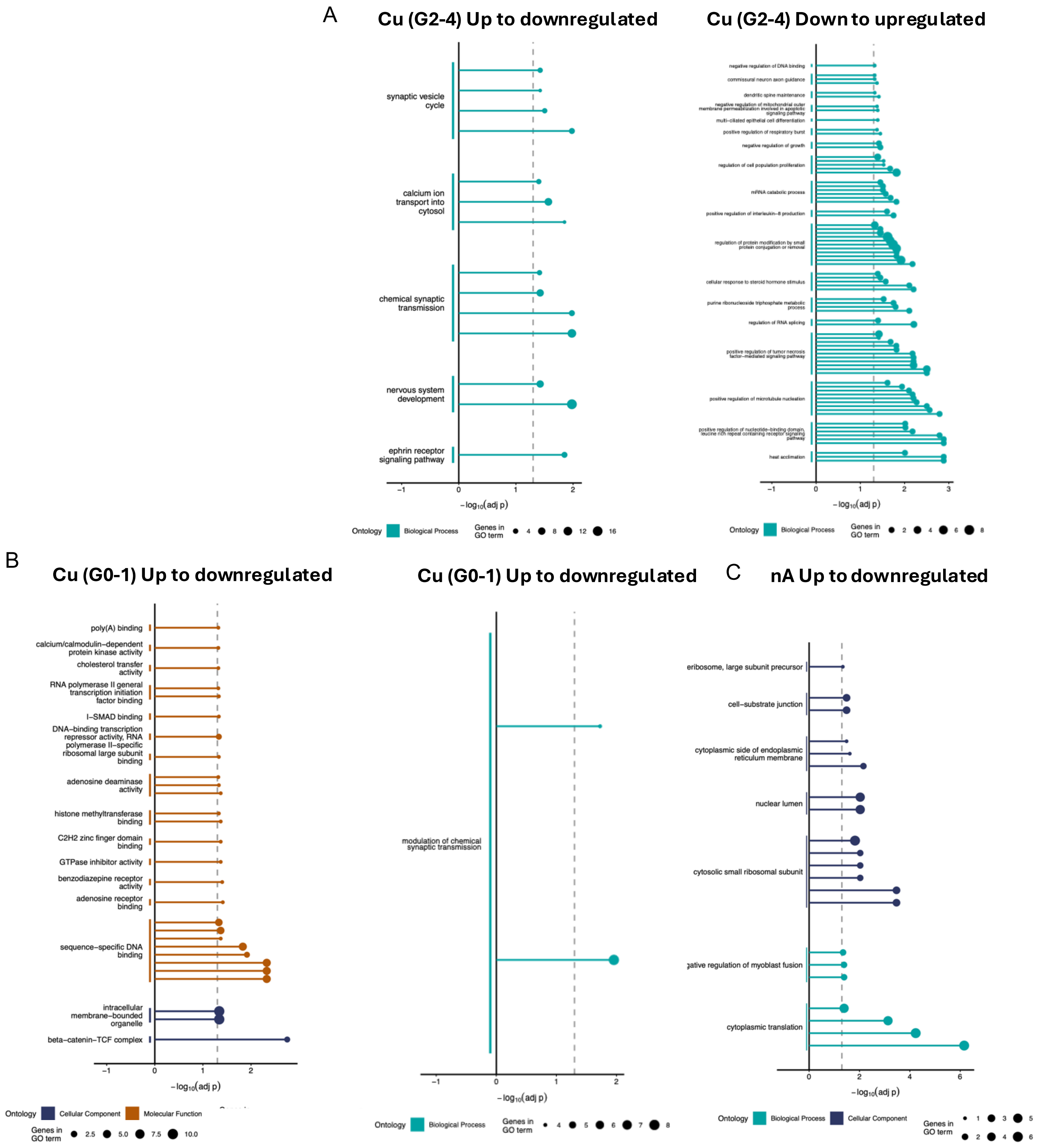


**Supplementary Figure 5: Lollipop plots of GEA changed direction DEGs between HD WGE and HD PM tissues.** Lollipop plots of GEA of the DEGs with opposing directions of change between WGE and adult HD PM tissue for ontologies BioP, MolF and CellC. The representative ontology term of each group is shown on the y axis. Each lollipop represents an ontology term: length represents −log10(adj p-value) of that term and size of the head represents the number of genes. Vertical dashed line represents p=0.05. **A** GEA of genes either up-to-down or down-to-up regulated in the Cu grades 2-4. **B** GEA of genes either up-to-down or down-to-up regulated in the Cu grades 0-1. **C** GEA of genes down-to-up regulated in the nA grades 2-4.

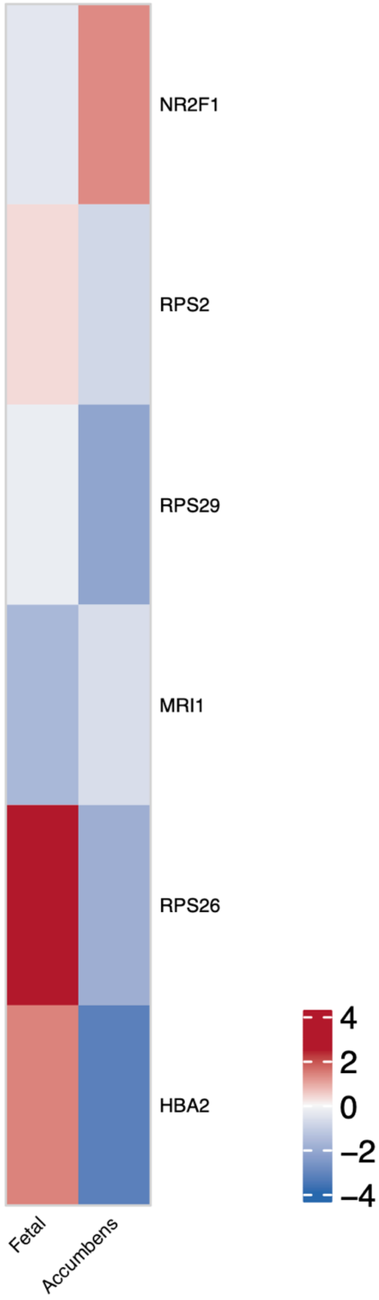


**Supplementary Figure 6: Heatmap of shared DEGs between HD WGE CGE cell types and nA PM.** Heatmap represents LogFC (red = upregulated, blue = downregulated; intensity represents magnitude)

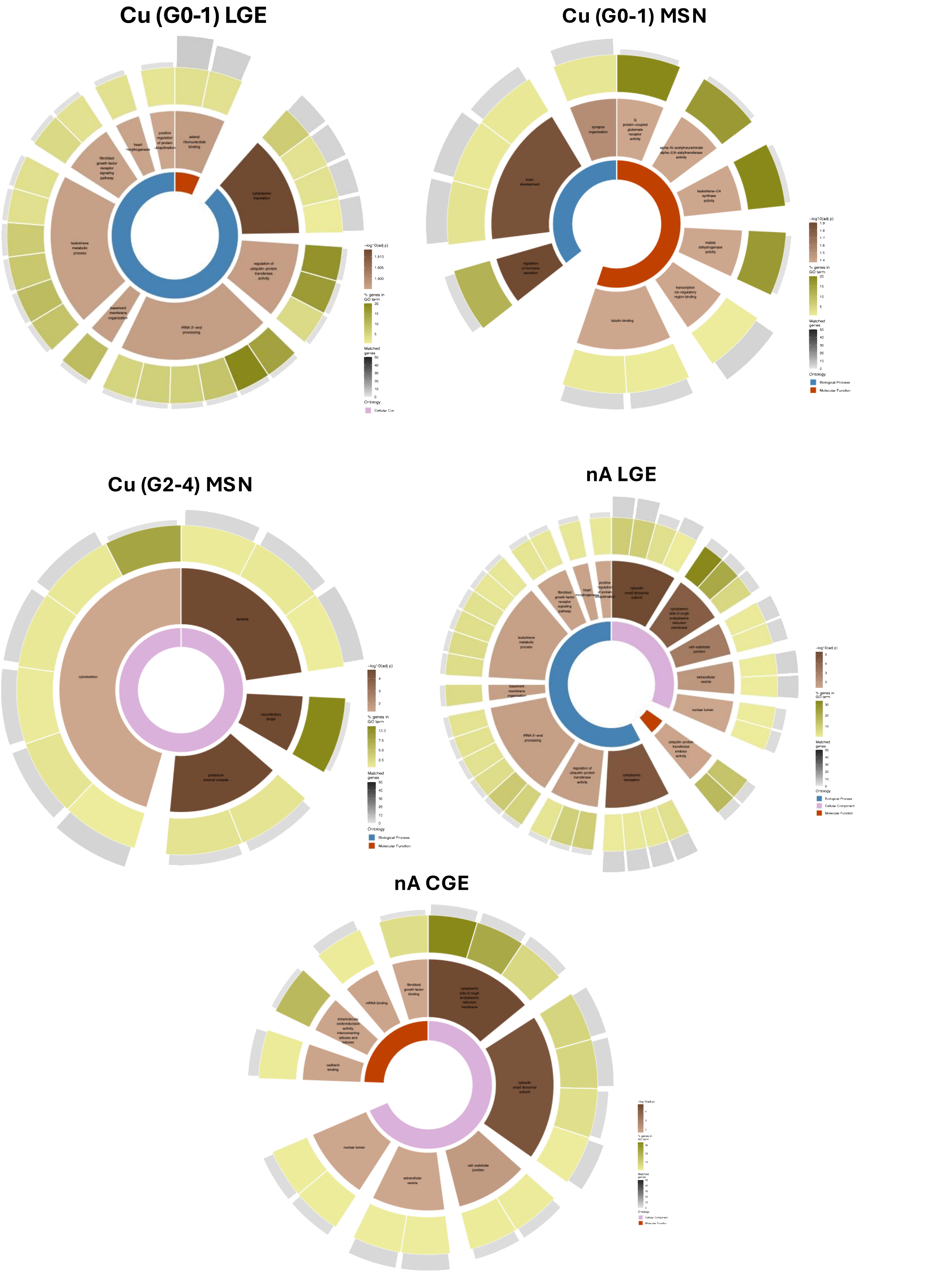


**Supplementary Figure 7: Sunburst plots from GEA on shared DEGs between HD LGE, MSN, and CGE clusters with HD PM Cu and nA tissues.** Inner ring: blue = BioP orange = MolF, pink = CellC; second ring (I-VI) represents parent categories, defined as the representative term of each group, coloured by −log10(adj p-value) of all GO terms within that group; third ring represents the DEGs as a % of the corresponding enriched term gene list (darker = higher %, paler = lower %); outer segments represent number of genes within an ontology term shown by length and colour. **A** GEA analysis of shared DEGs from LGE cluster and Cu grades 0-1, **B** GEA analysis of shared DEGs from MSN cluster and Cu grades 0-1, **C** GEA analysis of shared DEGs from MSN cluster and Cu grades 2-4, **D** GEA analysis of shared DEGs from LGE cluster and nA grades 2-4, **E** GEA analysis of shared DEGs from CGE cluster and nA grades 2-4,
